# Spontaneous network activity <35 Hz accounts for variability in stimulus-induced gamma responses

**DOI:** 10.1101/381236

## Abstract

Gamma activity is thought to serve several cognitive processes, including attention and memory. Even for the simplest stimulus, the occurrence of gamma activity is highly variable, both within and between individuals. The sources of this variability are largely unknown. They are, however, critical to deepen our understanding of the relation between gamma activity and behavior.

In this paper, we address one possible cause of this variability: the cross-frequency influence of spontaneous, whole-brain network activity on visual stimulus processing. By applying Hidden Markov modelling to MEG data, we reveal that the trial-averaged gamma response to a moving grating depends on the individual network profile, inferred from slower brain activity (<35 Hz) in the absence of stimulation (resting-state and task baseline). In addition, we demonstrate that dynamic modulations of this network activity in task baseline bias the gamma response on the level of trials.

In summary, our results reveal a cross-frequency and cross-session association between gamma responses induced by visual stimulation and spontaneous network activity.

## Introduction

Narrow-band gamma activity can be observed in numerous species and brain areas with various recording techniques (Bosman et al., 2014), including M/EEG recordings in humans (Jensen et al., 2007). It has been proposed to play a role in a variety of cognitive processes, including attention (Bosman et al., 2012; Grothe et al., 2012), feature binding (Engel et al., 1991; Singer and Gray, 1995), memory encoding (Sederberg et al., 2003; Jutras et al., 2009), memory retrieval (Osipova et al., 2006; Montgomery and Buzsaki, 2007), decision-making (van Wingerden et al., 2010, 2014), and reward processing (Berke, 2009; Kalenscher et al., 2010).

Importantly, gamma responses to visual stimuli vary substantially within and between subjects. Invasive recordings in monkeys (Lundqvist et al., 2016) and humans (Kucewicz et al., 2014) revealed that gamma responses of the same individual vary markedly from trial to trial. In fact, single-trial gamma responses have been described as transient events of varying amplitude, duration and frequency. These findings suggest that the oscillation-like appearance of the trial-averaged gamma response might be a misleading consequence of averaging, not reflecting the actual physiological processes engaged in single trials (Jones, 2016; van Ede et al., 2018). Still, averaging across trials results in a remarkably reproducible pattern, as shown by MEG studies measuring trial-average gamma responses in human visual cortex repeatedly in the same subjects (Hoogenboom et al., 2006; Muthukumaraswamy et al., 2010). Between subjects, in contrast, the trial-average response differs markedly with respect to amplitude, frequency and bandwidth (Muthukumaraswamy et al., 2010) and this between-subject variability has been shown to have a relatively strong genetic basis (van Pelt et al., 2012).

To date, the cause of within- and between-subject variability in gamma activity is not completely understood. Here, we propose that gamma responses might differ between subjects because subjects differ in their basic network profile underlying all brain activity. According to our hypothesis, these inter-individual differences become apparent even in the absence of gamma-inducing stimuli, implying that resting-state activity can predict gamma responses. This idea is based on functional magnetic resonance imaging (fMRI) (Smith et al., 2009; Cole et al., 2014, 2016; Tavor et al., 2016) and one recent MEG study (Becker et al., 2018), which demonstrated that resting-state network activity predicts inter-individual differences in task-related brain activity. Rest-task cross-frequency relationships affecting gamma oscillations, however, have not been investigated so far.

Notwithstanding the existence of robust network profiles, the brain is able to adapt flexibly to changes in the environment. Hence, we propose that network profiles are dynamic in nature and not only reflect the individual brain architecture, but also, possibly to a lesser extent (Gratton et al., 2018), the current situation. With respect to gamma activity, this assumption implies that induced responses might differ between trials because the individual network profile is modulated dynamically within a task.

To test these hypotheses, we derived an estimate of the individual network profile and its dynamics by applying Hidden Markov Modelling (HMM) to whole-brain MEG data, describing re-occurring patterns of network activity as repeated visits to a finite set of brain states (Fig. 1). Using the HMM, we investigated whether gamma responses differ between subjects because some subjects spend more time in certain brain states than others (between-subject effect). In addition, we tested whether the amplitude of the gamma response differs between trials because the pre-stimulus brain state differs between trials (within-subject effect). And finally, we compared the predictive potential of task-baseline vs. resting-state activity with respect to gamma amplitude.

**Fig. 1:**
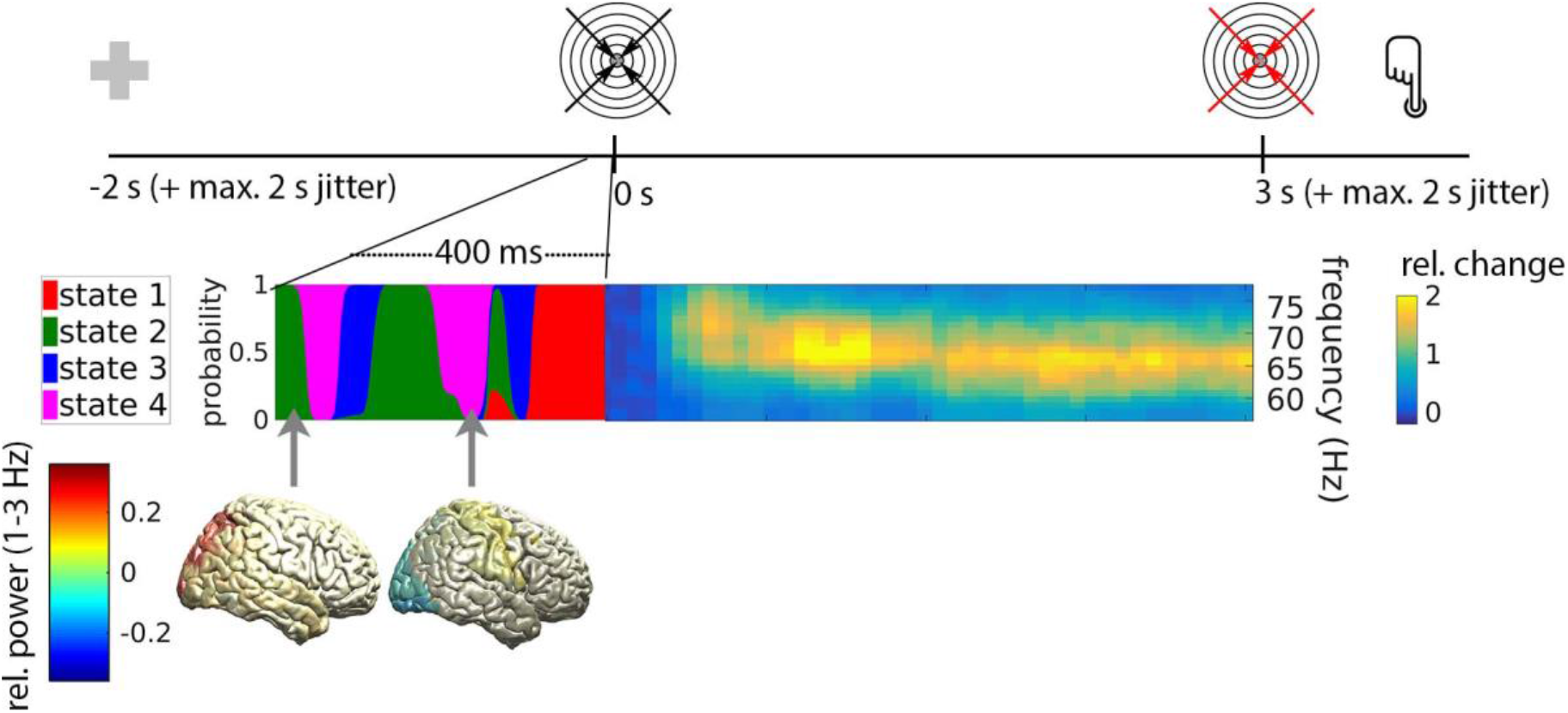
Experimental paradigm and rationale of the study. Upper row: Timeline of a single trial, locked to grating onset. Following a baseline period of 2 to 4 s with a central fixation cross, an inward-moving grating appeared which accelerated at an unpredictable moment 3 to 5 s following grating onset (illustrated here with red arrows). Subjects indicated they detected the acceleration via button press. Lower left: Hidden Markov Modelling yielded at each time sample the probability of the brain being in any of four states (color-coded). State inference was limited to epochs without stimulation, i.e. to the pre-stimulus baseline, as shown here, or to the resting-state (not shown). Each state is characterized by a unique spatio-spectral profile within the frequency ranges slower than gamma (1-35 Hz), including the topography of delta power shown here for states 2 (left) and 4 (right). Lower right: The inward-moving grating induced strong gamma activity in occipital areas. We investigated whether the strength of this stimulus-induced gamma response is related to spontaneously occurring whole-brain states.

## Methods

### Experimental Design

#### Participants

15 healthy participants were recruited for this study (21-45 years; 5 female). The study was approved by the Montreal Neurological Institute’s ethics committee (NEU 011-036) and was in accordance with the Declaration of Helsinki. All participants gave written informed consent and were compensated for their participation.

#### Paradigm

Subjects were presented with a modified version of the visual stimulation paradigm by Hoogenboom et al. (Hoogenboom et al., 2006): An inward-moving, circular sine wave grating with a diameter of 5° accelerated from 1.6 deg/s to 2.2 deg/s at an unpredictable moment between 3-5 seconds after stimulus onset. Subjects indicated that they had detected the velocity change by pressing a button with the index finger of the dominant hand. The button press ended the trial and the stimulus was turned off. During the inter-trial interval (baseline period), subjects were presented with a central fixation cross. Inter-trial intervals varied between 2 and 4s. A few trials with longer interval (17 - 19 s) were randomly interspersed in the trial sequence for all subjects but P1 (6 - 16 per subject; mean: 13). This was done to facilitate an analysis of the influence of baseline duration, but is not relevant for the analyses reported here.

#### Experimental Procedure

Each session started with a 5 min resting-state recording with eyes open, which was immediately followed by task practice and task recording. Before the start of the reaction time task, participants completed 10 practice trials. The task was divided into 2 - 5 blocks, containing 35 - 78 trials each (mean: 62.85). After each block, participants received a feedback on the accuracy of their responses and had the possibility to take a break. Following the reaction time task, a further 5 min resting-state recording was acquired.

In two subjects (P3 and S006R), additional task data were acquired 6 days and 1 day after the first recording session, respectively. Subject P1 was not recorded in resting state.

### Data acquisition

Participants were measured in a seated position with a 275-channel VSM/CTF MEG system at a sampling rate of 2400 Hz (no high-pass filter, 660 Hz anti-aliasing low-pass filter). Electrocardiography (ECG) and vertical electrooculography (EOG) were recorded simultaneously using MEG-compatible electrodes. Magnetic shielding was provided by a magnetically-shielded room with full 3-layer passive shielding. Participant preparation consisted of affixing 3 head-positioning coils to the nasion and both pre-auricular points. The position of the coils relative to the participant’s head was measured using a 3-D digitizer system (Polhemus Isotrack, Colchester, USA).

A T1-weighted MRI of the brain (1.5 T, 240 x 240 mm field of view, 1 mm isotropic, sagittal orientation) was obtained from each participant either at least one month before or immediately after the session.

### Preprocessing

Data were preprocessed and analysed using the HMM-MAR (Vidaurre et al., 2016) and Fieldtrip (Oostenveld et al., 2011) toolboxes for Matlab (The Mathworks). All data were screened visually. Noisy channels and noisy epochs were excluded from analysis. Data were down-sampled to 250 Hz. A 60 Hz discrete Fourier transform filter was applied to remove line noise. Cardiac and eye movement artefacts were isolated by FASTICA (Hyvarinen, 1999) and removed in non-automatic component selection.

### Source reconstruction

Individual T1-weighted MR scans were aligned to the MEG’s coordinate system, segmented and used for the construction of a single-shell, realistic head model (Nolte, 2003). To define a set of source coordinates, the “colin27” template MRI (Holmes et al., 1998) was inflated using FreeSurfer (Fischl et al., 1999) and a cortical mesh consisting of 2052 sources was constructed using MNE (Gramfort et al., 2014). The corresponding coordinates in individual head space were obtained by applying the inverse of the normalizing transform matching the individual to the template MR scan. The lead field (forward model) was computed based on the source coordinates and the head model. Subsequently, a Linearly Constrained Minimum Variance (LCMV) spatial filter (Van Veen and Buckley, 1988) was computed based on the lead field and the sensor covariance matrix, and data were projected through this filter trial-by-trial. Note that the sign of the beamformer output is arbitrary. Because our analysis requires sign consistency across subjects, we applied a sign flipping procedure to maximize sign consistency; see (Vidaurre et al., 2016) and (Vidaurre et al., 2018b) for details.

To reduce dimensionality, we grouped sources into parcels, defined by the Talairach Tournoux atlas (52), and carried out all subsequent analyses on the parcel level. First, each source was either assigned to one of 25 bilateral brain areas of interest or discarded from further analysis if it was more than 5 mm away from an area of interest (325 of 2052 sources). The areas of interest consisted of all cortical areas contained in the atlas, with the exception of seven areas at the base of the brain or deep within the interhemispheric fissure, which were assumed to have poor MEG signal quality (rectal gyrus, parahippocampal gyrus, subcallosal gyrus, transverse temporal gyrus, orbital gyrus, and uncus). The 25 bilateral brain areas of interest were further sub-divided into a left- and a right-hemispheric parcel, resulting in 50 cortical parcels of interest (Tab. 1 of the Supplementary Material). The first principle component was extracted from each parcel of interest and magnetic field spread between parcels was reduced by symmetric, multivariate orthogonalization (Colclough et al., 2015).

### Stimulus-induced gamma activity

We quantified post-stimulus gamma responses in order to relate them to brain states inferred from slower activity (≤35 Hz) occurring in the pre-stimulus baseline period or the resting-state recordings. Post-stimulus gamma responses were computed by multitaper spectral estimation using 2 Slepian tapers (Thomson, 1982). Power was estimated for frequencies between 40 Hz and 100 Hz in a 300 ms sliding window which was moved in steps of 50 ms. We screened post-stimulus parcel activity and identified a frequency band, a time window and a location of interest. Because individual gamma peak frequencies varied markedly across subjects (between 42 and 74 Hz), frequency selection was subject-specific, i.e. we defined an individual gamma band for each subject (individual gamma peak frequency ±10 Hz). The time window of interest was set to 0.6 to 2 s relative to stimulus onset because all subjects were found to exhibit stable gamma activity in this window. The bilateral cunei were chosen as the locations of interest because this was generally the area with the strongest gamma response. For the analyses described in the following, gamma power within ±10 Hz of individual gamma peak frequency was normalized frequency-by-frequency by computing the percent change relative to mean power in the response baseline (−0.5 to −0.2 ms from grating onset) and averaged over frequency, time and locations of interest.

### Hidden Markov Models

HMMs are probabilistic sequence models that find recurring patterns in time series data (Rabiner and Juang, 1986). Unlike sliding-window approaches, they can reveal fast state changes present in multichannel, electrophysiological recordings (Baker et al., 2014; Vidaurre et al., 2016, 2018a, 2018b). HMMs describe the dynamics of brain activity as a sequence of transient events, each of which corresponds to a visit to a particular brain state. For each state, the HMM infers a time-course that describes the probability of that state being active. Furthermore, each state is characterized by a unique spatio-spectral profile. In summary, HMM brain states can be considered a compact description of multi-faceted, recurring patterns in dynamic network activity. HMMs have been widely used in a variety of applications, such as the decoding of speech (Varga and Moore, 1990), the comparison of nucleotide sequences (Eddy, 1998) or the detection of pathological brain signals (Hirschmann et al., 2017; Kottaram et al., 2019).

### State inference

States were inferred separately from the baseline periods of the task, the rest recording preceding the task, and the rest recording following the task. For the baseline period, the first second of each trial was removed because it was assumed to contain activity related to the button press of the previous trial. Next, we *z*-scored and concatenated the data from all subjects in time, resulting in a total of 71.44 min of pre-task rest data (per subject mean: 5.10 min, STD: 1.40 min), 164.85 min of baseline data (per subject mean: 10.99 min, STD: 2.19 min) and 63.47 min of post-task rest data (per subject mean: 4.53 min, STD: 0.69 min). Importantly, we applied a spectral filter with a pass-band of 1 to 35 Hz to ensure that brain states were not based on gamma activity. This was done to demonstrate the universality of rest-task/baseline-task interactions, which we hypothesized to occur across frequency bands.

State inference was performed by applying a variety of the HMM designed to capture transient patterns of power and phase-coupling, referred to as Time-delay Embedded HMM (TDE-HMM; (Vidaurre et al., 2018b). In this model, each state is characterized by certain patterns of cross-correlation, which contain spectrally-defined patterns of power and phase-coupling. The TDE-HMM parameters were chosen as in (Vidaurre et al., 2018b).

Similar to the frequency resolution in spectral analysis, the number of states *K* in a HMM determines the level of detail of the solution. Here, we set *K*=4 to guarantee a reasonable amount of trials per state, and to provide enough level of detail to investigate the question at hand. Similar results were obtained for *K*=3.

### State properties

Following state inference, we computed the power and coherence associated with each state as detailed in (Vidaurre et al., 2016). In short, we used a Fourier-based multitaper approach to find the spectral properties of each state, restricting the estimation to the time points when a state was active. As a result, we obtained a multi-region pattern of power and coherence per state. For topographic illustrations, power and coherence were interpolated on a 3D-reconstruction of the template brain after computing the relative difference with respect to the mean over states. In case of coherence, we display the average coupling with all other parcels.

State dynamics can be summarized by statistics such as fractional occupancy (FO), lifetime and interval time (See Baker et al., 2014 for formal definitions). FO quantifies the fraction of samples assigned to a given state. Lifetime quantifies the duration of a state visit. Interval time quantifies the time in between subsequent visits of the same state.

### Statistical Analyses

#### Analysis of between-subject variability

We assessed whether the amplitude of trial-averaged, stimulus-induced gamma responses is related to state preferences in the baseline periods of the task and/or the rest recordings preceding and following the task. State probabilities were averaged across the entire baseline/rest recording and tested for a linear correlation with the amplitude of the trial-averaged gamma response using a significance test for Pearson’s correlation coefficient. Note that a strong correlation with any of the states will induce a correlation of opposite sign with the remaining states given the mutual exclusivity of states.

#### Analysis of within-subject variability

We hypothesized that the strength of stimulus-induced gamma activity depends on the brain state immediately before stimulus presentation (Fig. 1). To test this, we first computed, for each state, the average state probability in the pre-stimulus time window of interest, which served as state-specific trial weight. The pre-stimulus time window of interest was defined as −106 to 0 ms because 106 ms was the average state lifetime in task baseline (Supplementary Material). Subsequently, we computed a weighted trial average for each state using the obtained weights. This procedure can be considered a weighted (soft-assigned), within-subject grouping of trials by pre-stimulus state. Next, we tested whether the resulting trial groups consistently differed in post-stimulus gamma amplitude across subjects. This was achieved by running a Friedman test, followed by post-hoc Wilcoxon rank sum tests. Note that trials were grouped by pre-stimulus state, not post-stimulus gamma amplitude, i.e. any consistent difference in gamma amplitude must be due to a relationship between pre-stimulus state and post-stimulus gamma amplitude.

#### Comparison of effect size

We compared the between-subject and the within-subject effects of network activity on gamma responses by correlating FO with the amplitude of the gamma response. To separate between-subject from within-subject effects for both FO and gamma activation, we regressed out the average value for each subject from the single trial time courses. Pearson correlation between the subject-specific averages yielded the between-subject effects, and the correlation between the residual, trial-specific values yielded the within-subject effect. Note that trial-specific values could only be obtained for baseline, not for resting-state recordings. FO was originally represented as a 2-dimensional matrix (number of trials x number of states). To obtain a single value per trial, we applied Principal Component Analysis and kept only the first principal component.

In order to assess the variability of the different effects, we generated 1000 bootstrapped samples for each effect, using random sampling with replacement on the level of subjects to produce an empirical distribution of correlations (Rindskopf, 1997). *p*-values were obtained by computing the fraction of random samples with correlation 1 > correlation 2.

## Results

### Between-subject effect of brain states

In this study, we described 1-35 Hz spontaneous network activity by applying an HMM to resting-state MEG recordings and to the baseline periods of a task, respectively. An HMM estimates, for each sample of multivariate data, the probability of belonging to each of *K* possible brain states.

Using this approach, we derived four brains states from the resting-state recording acquired before the task (rest-pre), the baseline periods within the task (BL), and the resting-state recording acquired after the task (rest-post), respectively. Whereas it is possible to describe the data using more states, four were adequate for our purposes (see Materials and Methods).

We analysed the spectral properties of these brain states for each recording separately. Fig. 2A shows the spatial distribution of state power in the delta (1-3 Hz), theta (4-7 Hz), alpha (8-12 Hz), and the beta band (13-30 Hz) for the rest-pre recording. Fig. 2B depicts the state-specific power spectra, averaged over brain areas. Fig. 2C shows the correlations between state probability and the amplitude of the trial-averaged, stimulus-induced gamma response. Fig. 3 and Fig. 4 provide the corresponding information for task baseline and rest-post, respectively. Figures on state coherence are provided in the Supplementary Material (Fig. S1-S3).

**Fig. 2:**
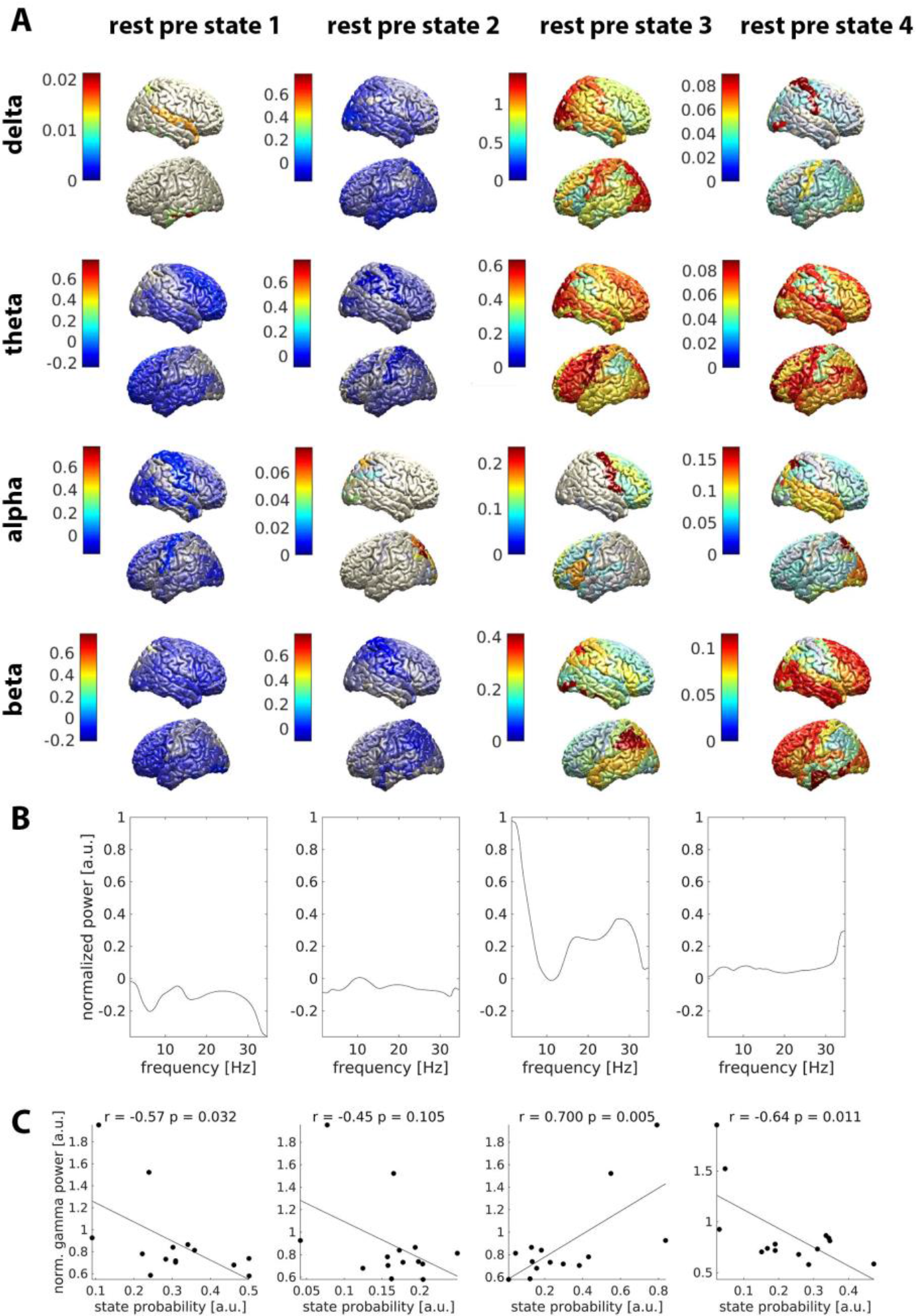
Brain states in the resting-state recording preceding the task. **A**: Topography of power for each state and frequency band. Relative difference to the mean over states is color-coded. **B**: Power averaged over parcels, normalized as in A. **C**: Correlation between state probability and the gamma response to visual stimulation. *r* = Pearson correlation. *p* = p-value.

**Fig. 3:**
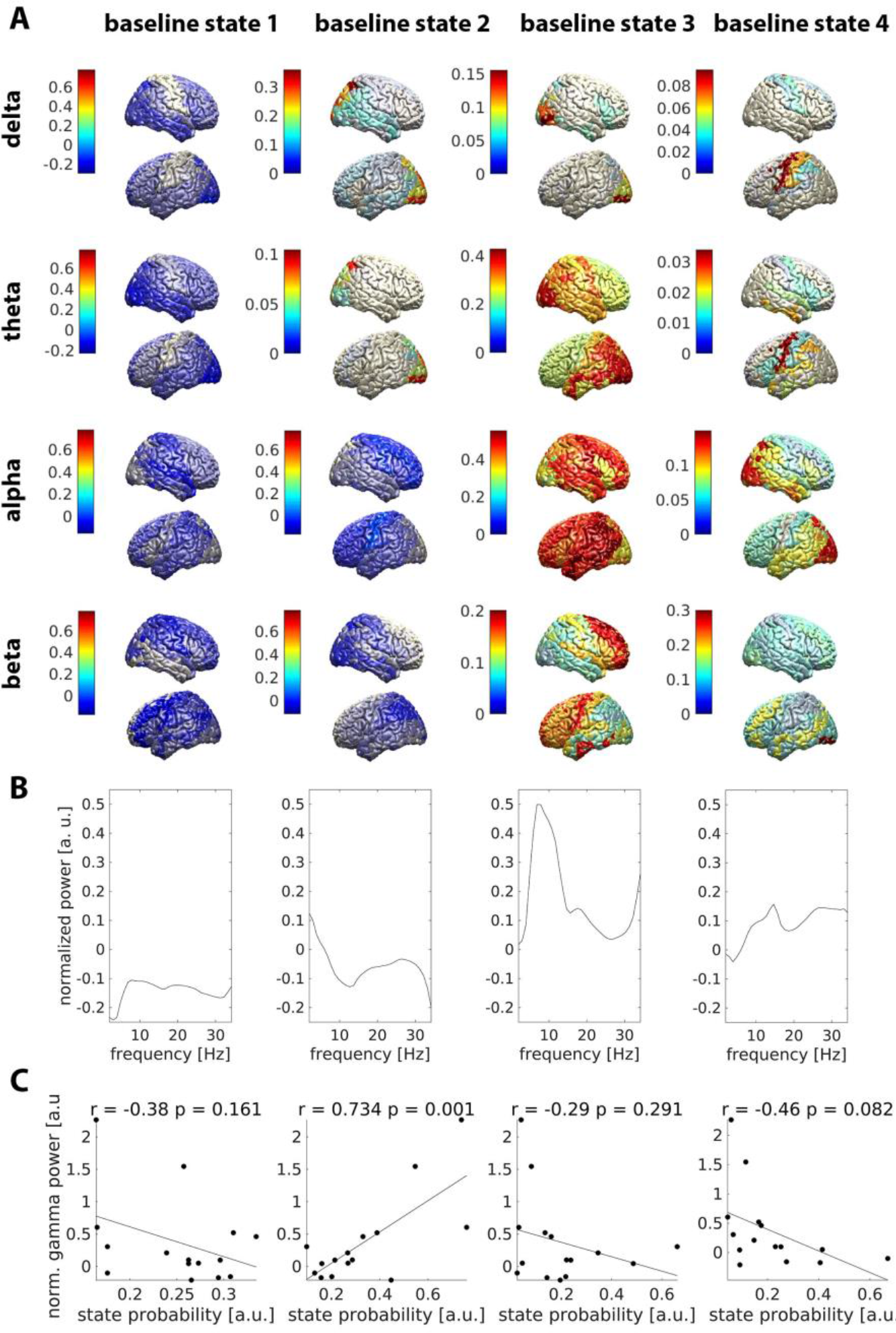
Brain states in the baseline periods of the task. **A**: Topography of power for each state and frequency band. Relative difference to the mean over states is color-coded. **B**: Power averaged over parcels, normalized as in A. **C**: Correlation between state probability and the gamma response to visual stimulation. *r* = Pearson correlation. *p* = p-value.

**Fig. 4:**
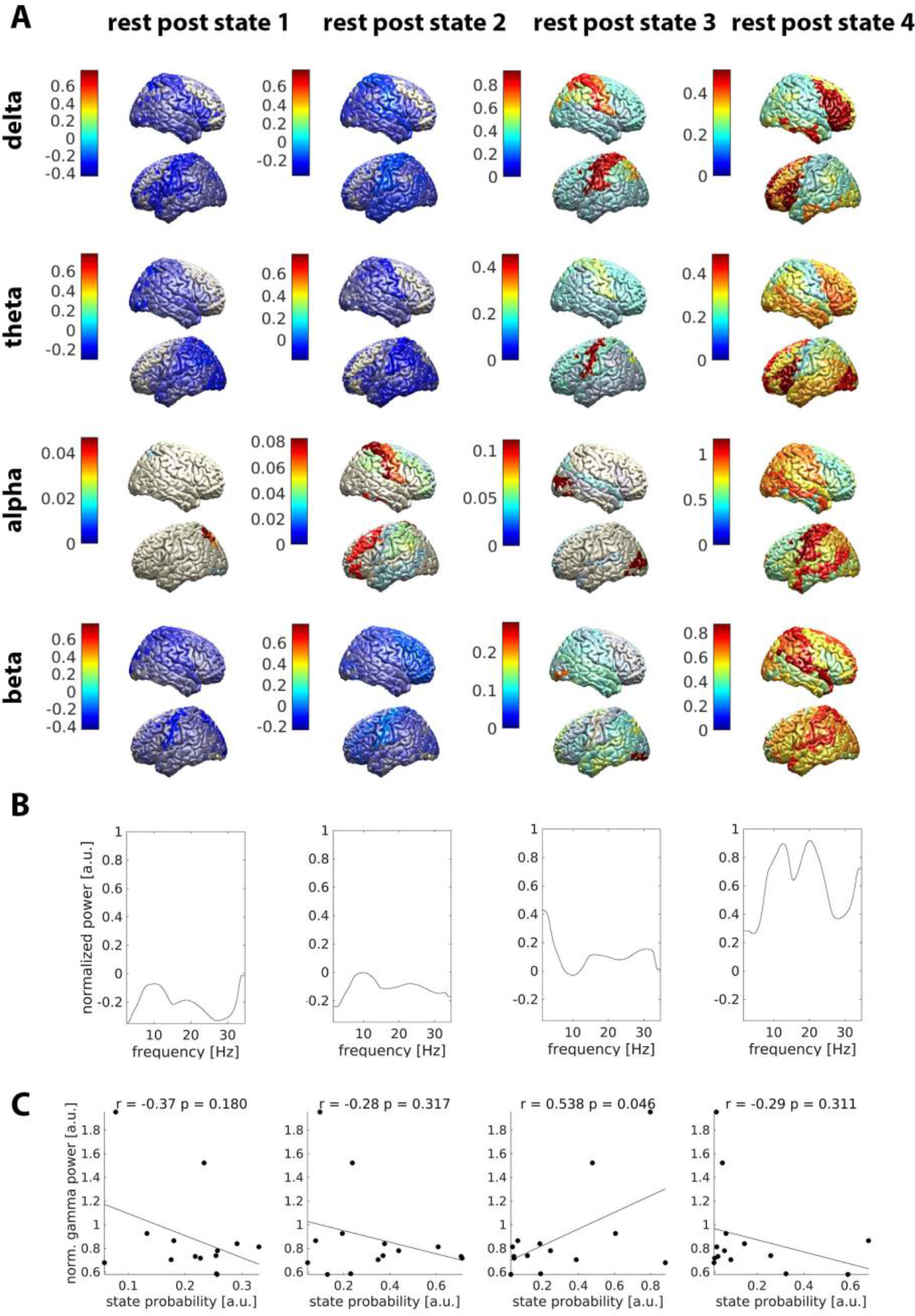
Brain states in the resting-state recording following the task. **A**: Topography of power for each state and frequency band. Relative difference to the mean over states is color-coded. **B**: Power averaged over parcels, normalized as in A. **C**: Correlation between state probability and the gamma response to visual stimulation. *r* = Pearson correlation. *p* = p-value.

We next focused on the brain state that was most strongly correlated with trial-average gamma amplitude (state 3 in rest-pre, state 2 in BL, state 3 in rest-post). The estimate of the individual preference for this state, defined as the fraction of time spent in this state, was similar for all recordings (Fig. S6 of the Supplementary Material). In all cases, this state showed a positive linear relationship with the gamma response, inducing a negative correlation with the remaining states due to the mutual exclusivity of states. The power spectrum of the positively correlated brain state was characterized by a peak in the delta/theta frequency range and a minimum in the alpha band (see Fig. 5A for a direct comparison of recordings). Delta power was concentrated in bilateral parietal and motor areas (Fig. 5B).

**Fig. 5:**
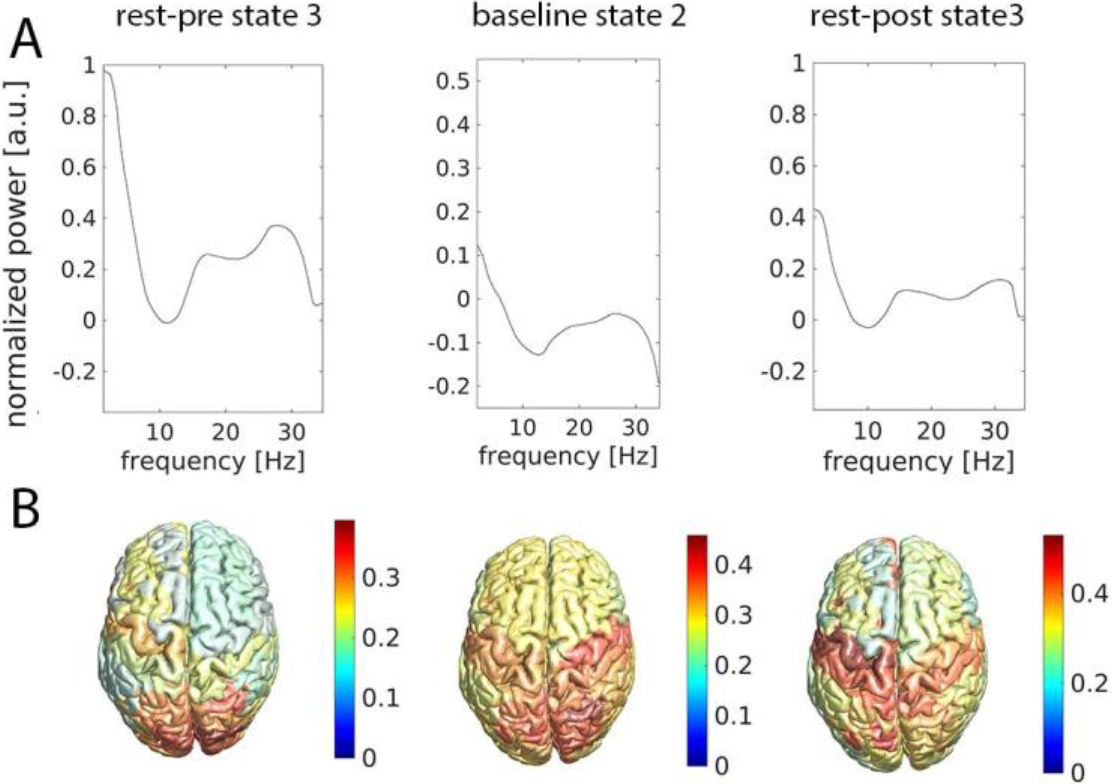
A common pattern found in all recordings. **A**: Normalized power spectra averaged over all parcels for rest-pre state 3 (left), baseline state 2 (middle) and rest-post state 3 (right). **B**: Spatial distribution of power between 1 and 5 Hz for the same states as in A. The relative difference to the mean over all rest-pre, baseline, and rest-post states is color-coded.

These findings demonstrate that (i) the HMM could identify a recurring pattern characterized by high delta and low alpha power in all of the recordings and (ii) that subjects showing this particular pattern frequently in the absence of a stimulus or task have a comparably strong gamma response.

### Within-subject effect of brain states

We tested whether the BL state occurring immediately before stimulus onset affects the amplitude of the gamma response within subjects. As illustrated in Fig. 6, gamma responses were strongest when BL state 2, i.e. the state positively correlated with the trial-average gamma response between subjects (see above), preceded grating onset. This pattern was observed in most individual subjects and was not an artefact of the trial-weighting procedure used in the analysis (Supplementary Material). In another control analysis, we verified that incompletely removed pre-stimulus eye blinks, which might impact the subsequent gamma response, were not the cause of pre-stimulus state changes (Fig. S4 of the Supplementary Material).

**Fig. 6:**
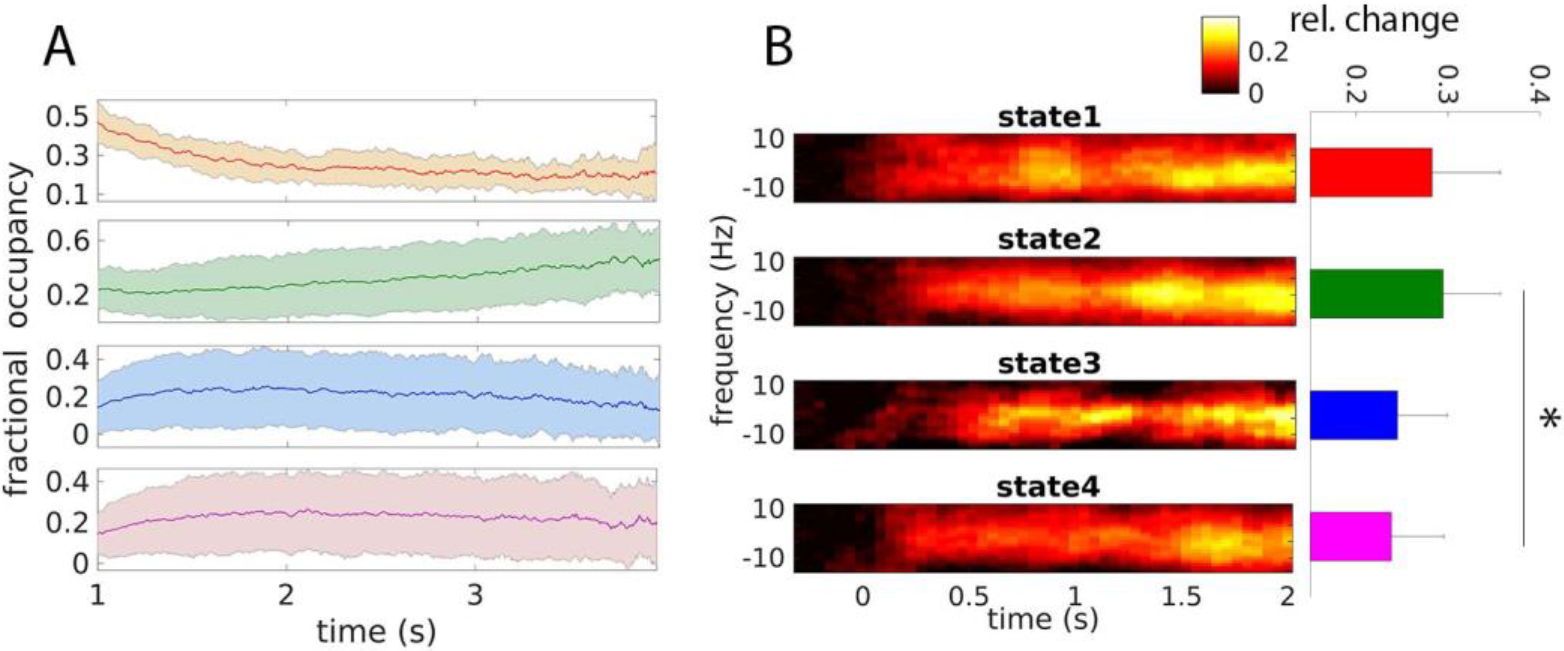
Relationship between gamma responses and pre-stimulus brain state. **A)** Fractional occupancy in the baseline period (1-4 s before grating onset), averaged over subjects and time-locked to the onset of the fixation cross. Shaded areas indicate the standard deviation over subjects. The first second of each trial was discarded to reduce the effect of movement-related processing occurring after the button press. **B)** Left: Weighted average time-frequency representations of gamma responses, time-locked to the appearance of the moving grating. 0 Hz marks individual gamma-peak frequency (between 42 and 74 Hz). Power was baseline-corrected (−0.5 to −0.2 s from grating onset). Right: Power averaged over frequency (individual gamma peak frequency ±10 Hz) and time (0.6 to 2s).

Due to the association with a strong gamma response both within and between subjects, we refer to BL state 2, which corresponds to rest-pre state 3 and rest-post state 3, as the “gamma-enhancing brain state” in the following.

We investigated whether the gamma-enhancing brain state was upregulated as the baseline period progressed, in anticipation of the stimulus. To this end, we averaged the state probabilities, time-locked to the beginning of the baseline period (fixation-cross onset), over trials. The resulting average, referred to as fractional occupancy (FO), quantifies how often a given state occurred at each time point in the baseline period (Baker et al., 2014). Indeed, BL state 2 appeared more frequently towards the end of the baseline period (Fig. 6; mean slope = 0.086, p < 0.001; *t*-test). BL state 1 was found to be predominant early in the baseline, but its FO decreased over time (mean slope = −0.065, p < 0.001; *t*-test). The FO of BL state 3 showed a weak negative dependency on time (mean slope = − 0.020, p = 0.02; *t*-test) and BL state 4’s FO did not change significantly (mean slope = −0.001, p = 0.87; *t*-test).

Finally, we considered the possibility that BL state 2 owes its gamma-enhancing properties solely to low pre-stimulus alpha power in parieto-occipital cortex. To explore this possibility, we split the trials in two groups based on alpha power observed in the cuneus over the last 100 ms before stimulus onset (lower 50% vs. higher 50%). Stimulus-induced gamma power did not differ between these groups (p = 0.43; Friedman test). Similar results were obtained when comparing three levels of alpha power and when considering all parieto-occipital brain areas instead of only the cuneus (data not shown). These results suggest that low parieto-occipital alpha power does not fully explain the within-subject effect detected here, highlighting the relevance of whole-brain spontaneous network dynamics.

### Comparison of effect size

So far, we have revealed four different effects of spontaneously occurring brain states on stimulus-induced gamma responses: an across-subject correlation for rest-pre, an across-subject correlation for task baseline, an across-subject correlation for rest-post, and a within-subject effect for task baseline. We now compare the strength of these different effects, finding a dominance of between-subject over within-subject effects. A quantitative comparison of effect size is displayed in Fig. 7. The within-subject effect was much weaker than any of the between-subject effects. Qualitatively, the between-subject effect was stronger for task baseline than for rest-pre and for rest-post, respectively. These differences, however, were not significant (BL vs. rest-pre: *p* = 0.25, BL vs. rest-post: *p* = 0.17).

**Fig. 7:**
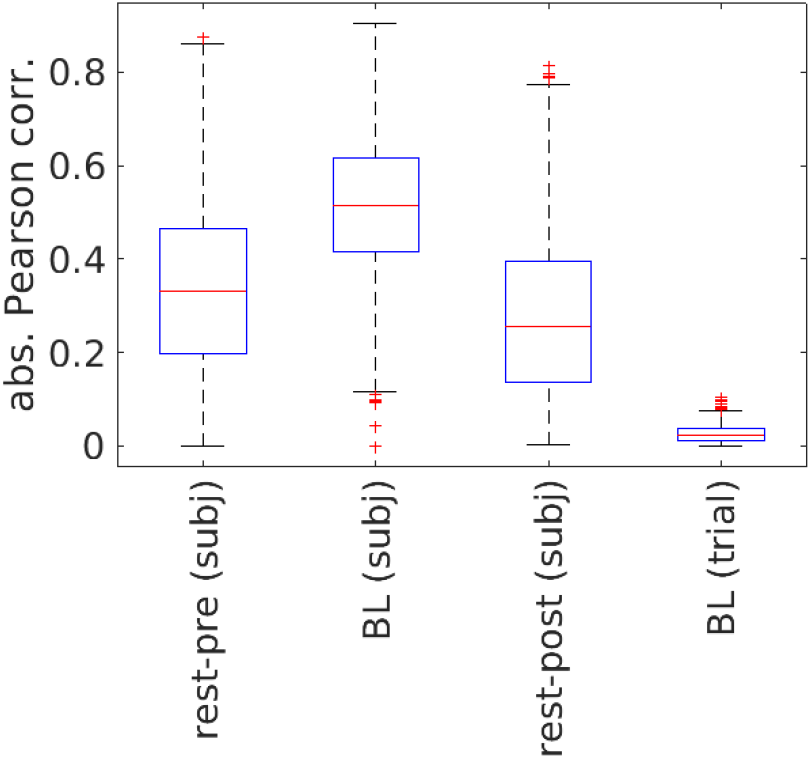
Comparison of effect size. Resampling subjects with replacement yielded empirical distributions of the absolute Pearson correlation coefficient. The distribution medians are represented by red lines and the 0.25 – 0.75 interquartile range (IR) is indicated by black whiskers. Outliers (median ± 2.5 IR) are represented by plus signs. *rest-pre (subj)*: between-subject correlation for resting-state recording acquired before the task; *BL (subj)*: between-subject correlation for baseline periods of the task; *rest-post (subj)*: between-subject correlation for resting-state recording acquired after the task. *BL (trial):* within-subject correlation for baseline periods of the task.

## Discussion

In this paper, we have demonstrated that inter-individual differences in gamma responses to visual stimulation are reflected by inter-individual differences in spontaneous network activity <35 Hz. Furthermore, we have revealed a similar, albeit weaker, influence of brain states on trial-specific gamma responses. Our results imply that it is possible to predict a subject’s gamma response from their resting-state activity profile.

### Hidden Markov Modelling of brain activity

The HMM has several useful properties for network-level analysis of electrophysiological data. Unlike sliding-window approaches, it processes the data sample by sample, facilitating the characterization of electrophysiological networks at very high temporal resolution. In addition, the HMM is a multivariate approach that considers all signals simultaneously. Rather than defining a region of interest, one can process all brain areas at once. Subsequent statistical tests do not need to be corrected for multiple comparisons, improving statistical efficiency. Importantly, the HMM is not a biophysical model explaining how brain activity arises, but a data-driven approach providing a compact representation of multi-channel/multi-area data.

Recent MEG studies made use of these properties to reveal a rich repertoire of fast-changing network states characterized by distinct topographies of spectral power and coupling, many of which were reminiscent of the resting-state networks originally obtained with fMRI (Baker et al., 2014; Vidaurre et al., 2016, 2018a, 2018b). HMMs and other whole-brain models have also been applied to describe dynamic connectivity in fMRI data (Cabral et al., 2017; Vidaurre et al., 2017b). Here, we have used this approach to assess the relationship between spontaneous network activity (resting-state and task baseline) and stimulus-induced gamma responses.

### Interactions between spontaneous and task-related brain activity

A number of fMRI studies have previously demonstrated interactions between resting-state and task-related activity (Northoff et al., 2010). Resting-state and task-related networks were found to be highly similar across a wide variety of tasks (Smith et al., 2009; Cole et al., 2014). Furthermore, stimulus-induced patterns of activation could be predicted from resting-state activity (Cole et al., 2016; Tavor et al., 2016). These findings suggest that task-related brain activity arises by relatively minor modulations of a basic network profile, which can be considered a neural signature or fingerprint, allowing for accurate identification of individual subjects (Finn et al., 2015).

The current MEG study is one the first to show that the above concept might be transferable from fMRI to neurophysiology. The fact that the individual preference for a particular brain state correlated with the individual gamma response implies that spontaneous brain activity measured in the absence of stimulation (rest or task baseline) is predictive of brain activity induced by a visual stimulus. This observation supports the concept of a robust, individual network profile.

So far, there is only one comparable piece of work from Becker et al, who likewise combined MEG and HMMs to predict electrophysiological responses to visual stimuli and own movements from resting-state activity (Becker et al., 2018). The current study differs from this paper in several ways. First, it investigates induced rather than evoked responses. Second, it assesses within-subject variability in addition to between-subject variability. Third, it predicts gamma band responses from low-frequency activity (cross-frequency analysis). And finally, it compares the predictive potential of resting-state and task baseline activity.

A unique insight resulting from this study is that rest-task relationships exist across frequency bands, as evidenced by an influence of spontaneous 1-35 Hz activity on stimulus-induced responses in the gamma band (>35 Hz). A possible explanation might be that fundamental brain functions like attention, which do impact brain responses and might be reflected by brain states, involve predominantly theta and alpha oscillations. This possibility is discussed in more detail below (see Brain States and Attention).

In addition, our study shows for the first time that induced gamma responses in human visual cortex are biased by pre-stimulus, spontaneous brain activity below 35 Hz. While this finding aligns with similar observations made for spiking (Tsodyks et al., 1999), evoked responses in local field potentials (Arieli et al., 1996; Kisley and Gerstein, 1999) and the BOLD signal (Fox et al., 2006), as well as perception (e.g. van Dijk et al., 2008; Busch et al., 2009; Baumgarten et al., 2015), it highlights one of the major advantages of our approach. The combination of MEG and HMM provides network activity resolved on a millisecond time scale, thus providing insights on the level of subjects *and* trials. On the one hand, the approach allows for estimating a subject’s average gamma response based on the brain states generally preferred by this subject. On the other hand, the same model allows for estimating the gamma response in the current trial based on the brain state last visited before stimulus onset. Interestingly, we found the subject-level estimates to be much more precise than the trial-level estimates, indicating that the brain states described here are more representative of the current individual than of the current trial.

Finally, we investigated whether task baseline activity is more predictive of brain responses than resting-state activity. The first thing to note is that both kinds of recordings contained the same basic pattern related to gamma responses (“the gamma-enhancing brain state”), indicating that prediction depends on how well the individual preference for this pattern can be estimated from a given recording. While a recent fMRI study suggests that task recordings might allow for a better discrimination between individual network profiles than resting-state recordings (Greene et al., 2018), we observed only qualitative differences when predicting gamma responses. This speaks for a limited influence of context on individual network profiles (compare Gratton et al., 2018).

### Brain states and attention

The current study did not attempt to quantify attention, and thus it cannot establish a direct link between attention and brain states. Nevertheless, there are several observations indicating that spontaneous switching between brain states in part reflects the dynamic modulation of attention. First, attention can enhance gamma responses in visual cortex, similar to the gamma-enhancing brain state observed here (Tallon-Baudry et al., 2005). Second, the gamma-enhancing brain state became more common towards the end of the baseline period, which might reflect an anticipatory upregulation of attention as stimulus presentation approached. Third, the gamma-enhancing brain state is characterized by low posterior alpha power, which is believed to reflect the current level of attention. This view is grounded in M/EEG studies showing that briefly presented visual stimuli are more likely to be perceived if posterior alpha oscillations are desynchronized (Hanslmayr et al., 2007; Dijk et al., 2008; Lange et al., 2013). When subjects are instructed to pay attention to one visual hemifield, alpha power increases in the ipsilateral hemisphere, probably to reduce the influence of distractors in the irrelevant hemifield (Mazaheri and Jensen, 2008; Treder et al., 2011; Horschig et al., 2015). Similar observations have been reported for other sensory modalities (Haegens et al., 2011; Mazaheri, 2014; Baumgarten et al., 2016), suggesting that alpha power might serve as a general mechanism for controlling whether or not a stimulus is noticed by the subject. Finally, recent MEG studies demonstrated a relationship between pre-stimulus alpha and post-stimulus gamma oscillations in visual (Popov et al., 2017) and somatosensory areas (Wittenberg et al., 2018).

Importantly, however, the gamma-enhancing brain state described here cannot be equated with low posterior alpha power. Differences in alpha power did not entirely account for the effects of brain states in this study, and high delta rather than low alpha power was the common feature shared across states correlating positively with the gamma response. Thus, low alpha power in occipital and parietal areas might be just one aspect of sensory gating.

Assuming that brain states indeed reflect the level of attention, it is possible to give a rather parsimonious interpretation of our findings. First, the correlation between rest-based brain states and the trial-average gamma response indicates that it is possible to identify subjects capable of maintaining a high level of attention based on resting-state activity. This identification works equally well or even better when basing the identification on task baseline activity. Similarly, the analysis of pre-stimulus brain states allows for identifying trials in which stimulus presentation coincides with a high level of attention.

### Limitations

While this work demonstrates a correlation between resting-state activity and brain responses, it is limited to the amplitude of induced gamma responses in the visual system. Other features such as frequency or latency were not assessed. And, unlike previous fMRI studies (Cole et al., 2016; Tavor et al., 2016) and the MEG study by Becker et al. (Becker et al., 2018), it did not probe the predictive power of resting-state activity by generating out-of-sample predictions.

### Conclusions

We have shown that brains states describing spontaneous network activity < 35 Hz are correlated with the amplitude of stimulus-induced gamma responses. Our findings suggest that each subject is characterized by an individual network profile predictive of brain responses.

