## Supplementary Material for "Spontaneous network activity <35 Hz accounts for variability in stimulus-induced gamma responses"

#### Between-subject effect of brain states

##### *State coherence*

Fig. S1A shows the spatial distribution of state coherence in the delta (1-3 Hz), theta (4-7 Hz), alpha (8-12 Hz), and the beta band (13-30 Hz) for the rest-pre recording. Fig. S1B depicts the state-specific coherence spectra. Fig. S1C shows the correlations between state probability and the amplitude of the trial-averaged, stimulus-induced gamma response. Fig. S2 and Fig. S3 provide the corresponding information for task baseline and rest-post, respectively.

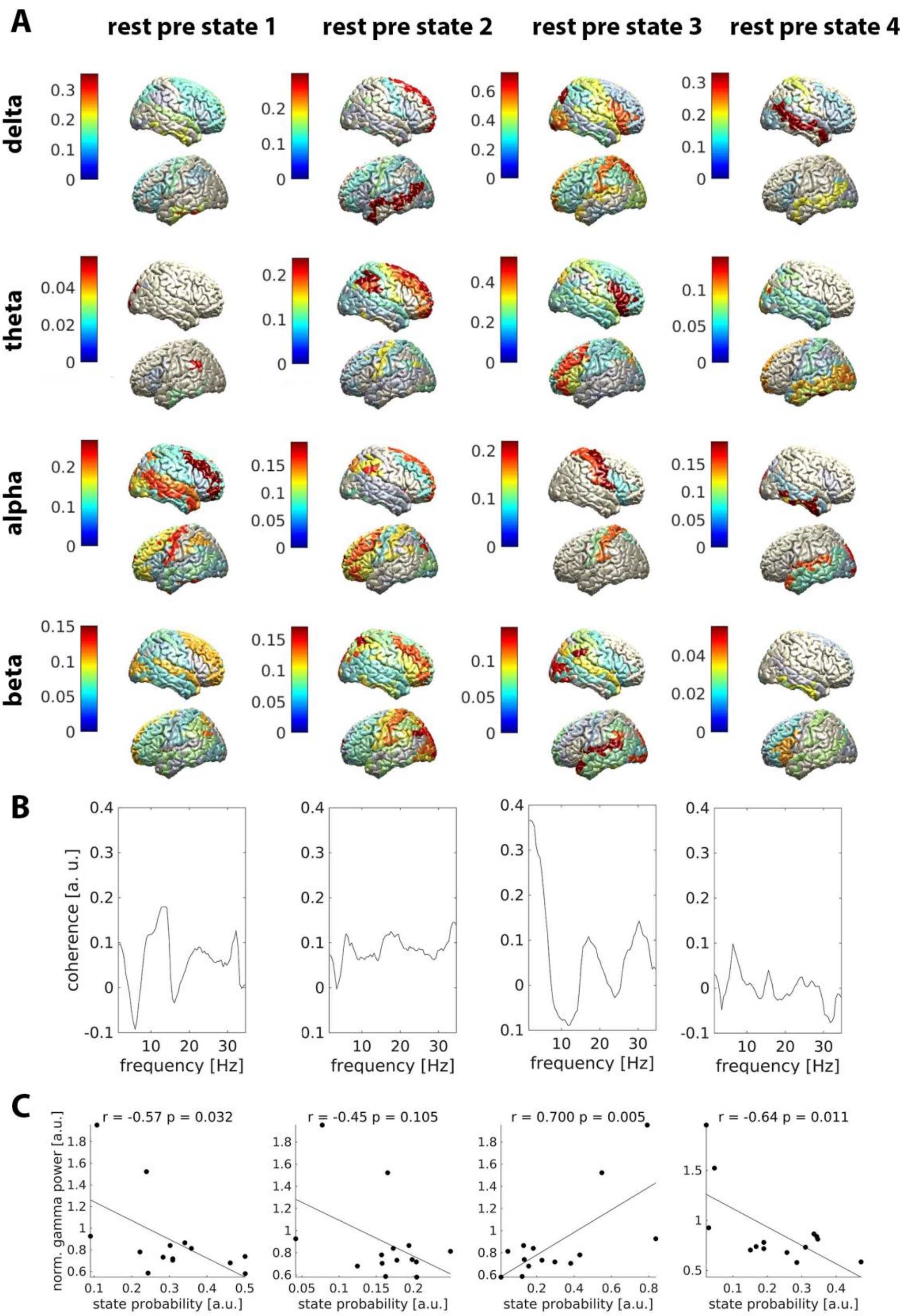


**Fig. S1:** *Brain states in the resting-state recording preceding the task*. **A**: Topography of coherence for each state and frequency band. Colours indicate the average coherence of each parcel with all other parcels, as relative difference to the mean across states. **B**: Coherence averaged over parcels, normalized as in A. **C**: Correlation between state probability and the gamma response to visual stimulation. *r* = Pearson correlation. *p* = p-value.

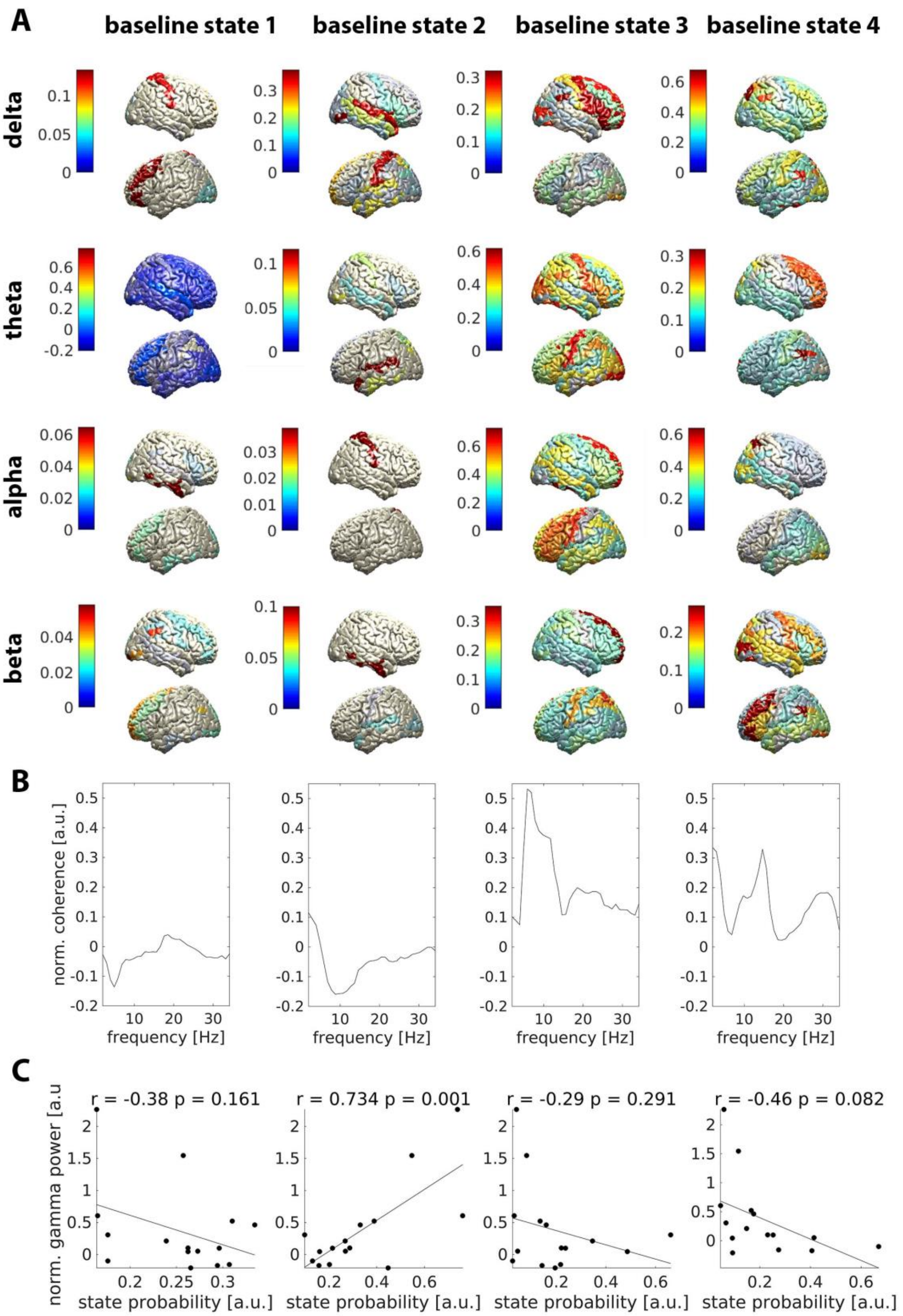


**Fig. S2:** *Brain states in baseline periods of the task*. **A**: Topography of coherence for each state and frequency band. Colours indicate the average coherence of each parcel with all other parcels, as relative difference to the mean across states. **B**: Coherence averaged over parcels, normalized as in A. **C**: Correlation between state probability and the gamma response to visual stimulation. *r* = Pearson correlation. *p* = p-value.

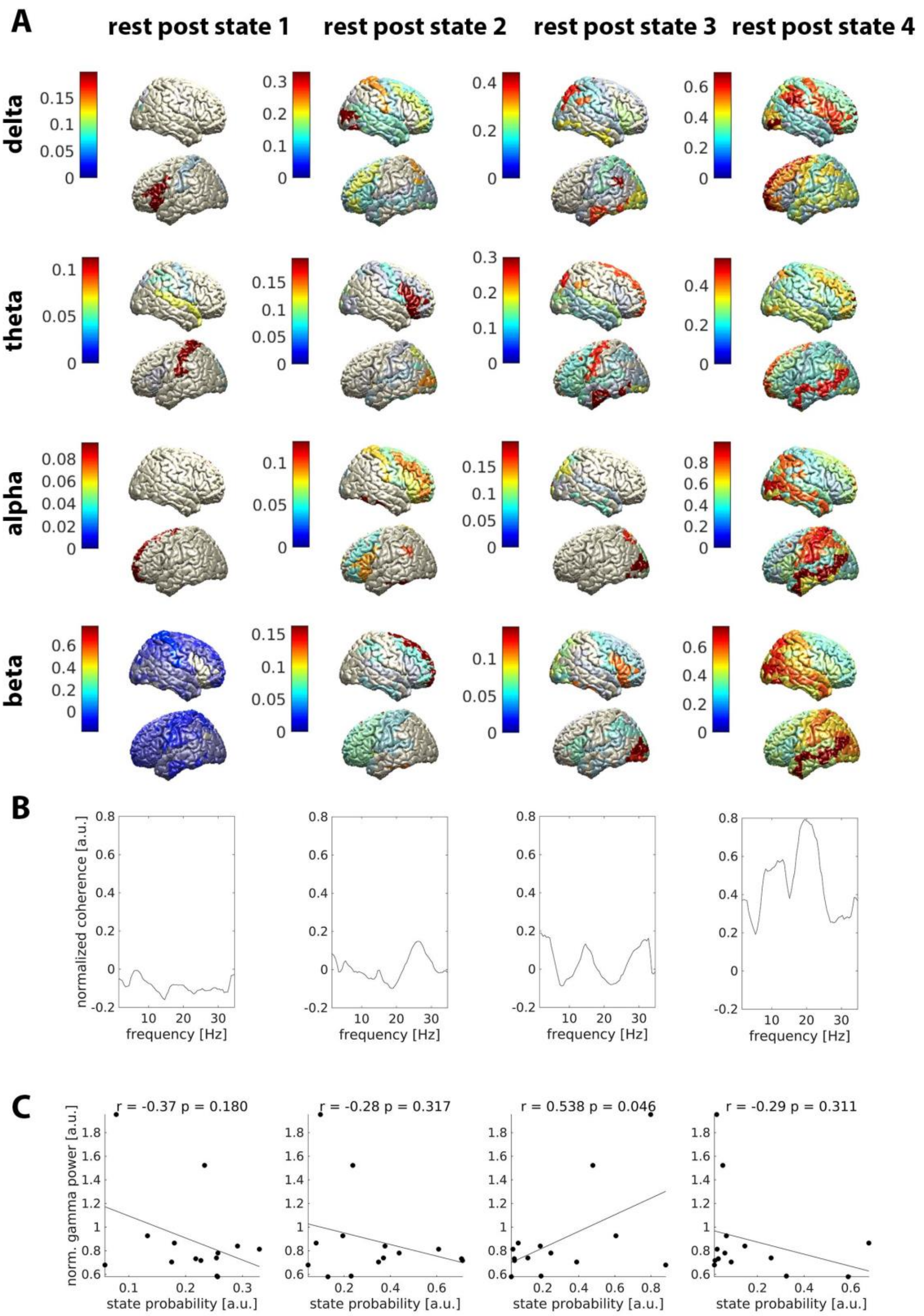


**Fig. S3:** *Brain states in the resting-state recording following the task*. **A**: Topography of coherence for each state and frequency band. Colours indicate the average coherence of each parcel with all other parcels, as relative difference to the mean across states. **B**: Coherence averaged over parcels, normalized as in A. **C**: Correlation between state probability and the gamma response to visual stimulation. *r* = Pearson correlation. *p* = p-value.

### Within-subject effect of brain states

Fig. S4 shows fractional occupancy, mean lifetime and mean interval time for each baseline state. Fractional occupancy quantifies the fraction of samples assigned to a given state. Lifetime quantifies the duration of a state. Interval time quantifies the time in between subsequent visits of the same state. See (1) for a formal definition.

Separate one-way ANOVAs revealed no difference in fractional occupancy ($F(3,56) = 0.76$, $p = 0.39$) and no difference in mean interval time ($F(3,56) = 0.32$, $p = 0.81$) between states. There was a difference, however, in mean lifetime ($F(3,56) = 6.51$, $p = 0.0007$). Tukey-Kramer corrected post-hoc tests showed that state 1 had a longer lifetime than states 3 (mean difference = 0.07, $p = 0.008$) and 4 (mean difference = 0.06, $p = 0.0063$). The mean lifetime over baseline states was 106 ms, which defined the pre-stimulus time window of interest for detecting pre-stimulus effects (see main text, Materials and Methods).

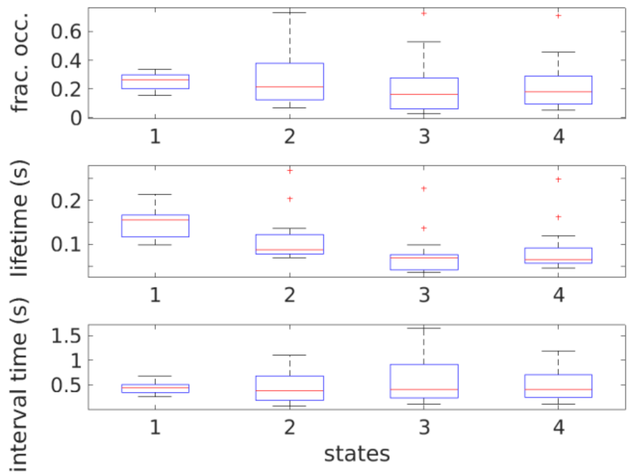


**Fig S4:** *Summary statistics of state occurrence in baseline epochs.* **Top**: Fractional occupancy. **Middle**: Mean lifetime. **Bottom**: Mean interval time.

To test whether baseline states depend on heartbeat or eye blinks, we detected events in the ECG and the EOG signal, respectively, by applying individually adjusted thresholds to the high-pass filtered (>3 Hz) and z-scored ECG/EOG data. Next, we investigated whether fractional occupancy changed around these events. As depicted in Fig. S5, neither heartbeat nor eye blinks were associated with changes in fractional occupancy, implying that neither heartbeat nor blinking caused systematic state changes.

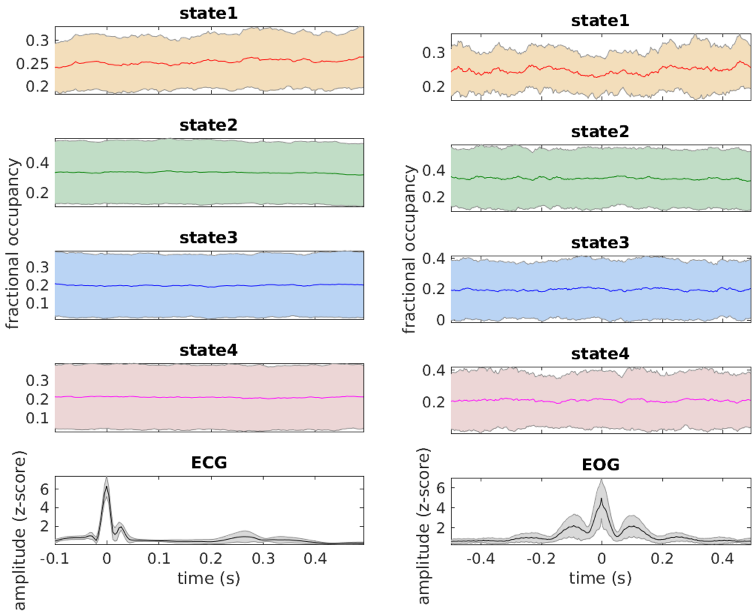


**Fig. S4**: *Fractional occupancy in relation to heartbeat (left) and eye blinks (right).* Shaded areas indicate the standard deviation over subjects. ECG = electrocardiogram, EOG = vertical electrooculogram.

#### *Association between pre-stimulus states and induced gamma activity in individual subjects*

Fig. S5 illustrates the relationship between pre-stimulus states and stimulus-induced gamma activity in individual subjects. 11 out of 15 subjects showed stronger gamma responses following baseline state 2 than following baseline state 4. The responses were approximately equal in three subjects (P1, S008 and S009) and the opposite pattern, i.e. stronger responses following baseline state 4, was observed in one subject (S007).

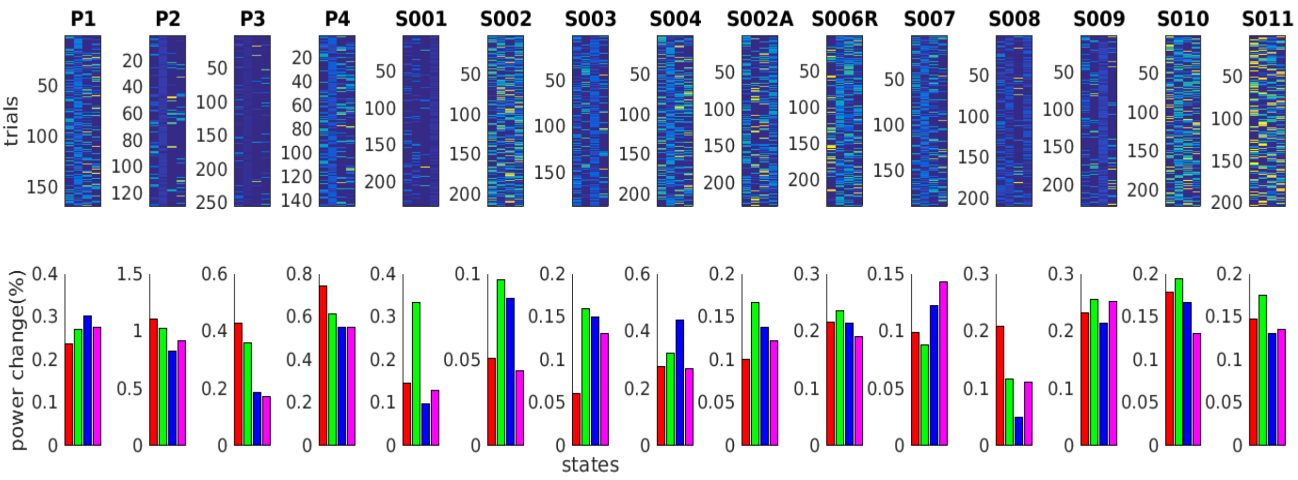


**Fig. S5**: *Relationship between pre-stimulus states and stimulus-induced gamma activity in individual subjects.* **Top:** Trial weights by state and subject (color-coded). **Bottom**: Amplitude of the stimulus-induced gamma response by preceding baseline state.

#### *Trial-weighting cannot explain the relationship between baseline states and stimulus-induced gamma activity*

In this paper, we compared weighted averages of stimulus-induced gamma power across brain states. The trial weights were based on the state probabilities in the pre-stimulus time window of interest. This trial-weighting procedure can produce weighted average time-frequency representations (TFRs) which are similar to single-trial TFRs, i.e. noisy, if a lot of weight is assigned to individual trials. As seen in Fig. S5, upper row, considerable weight was indeed assigned to individual trials in some subjects, such as P2. If the weight concentration is systematically different between states, this can result in systematic differences between state-specific TFRs, i.e. the weight concentration might confound the interpretation. To exclude that this was the case in our data, we quantified weight concentration by information entropy and compared entropy across states. Information entropy is defined as

$$H(X) = -\sum_{i=1}^{N} P(x_i) log P(x_i)$$

and describes the uncertainty about an outcome. It is maximal when all possible events have equal probability, i.e. when weight concentration is minimal. Importantly, we did not find a difference in entropy ($p$ = 0.26, Friedman test), suggesting that the differences between states are most likely not due the weighting procedure.

#### *Robustness of the individual preference for the gamma-enhancing network state*

The individual preference for the gamma-enhancing network state, indexed by the fraction of time spent in this state, was stable across recordings sessions, especially when comparing the resting-state recording preceding the task to the baseline periods of the task. Fig. S6A shows the linear correlations between the preference estimates obtained from different sessions (baseline vs. rest-pre: $r = 0.85$, $p < 0.001$; baseline vs. rest-post: $r = 0.53$, $p = 0.05$; rest-pre vs. rest-post: $r = 0.64$, $p = 0.01$). Fig. S6B depicts the ordering of subjects with respect to the probability of visiting the gamma-enhancing rest state. With some exceptions, subjects held a similar rank in each recording. These results suggest that the individual preference for the gamma-enhancing network state is a temporally stable characteristic of each subject.

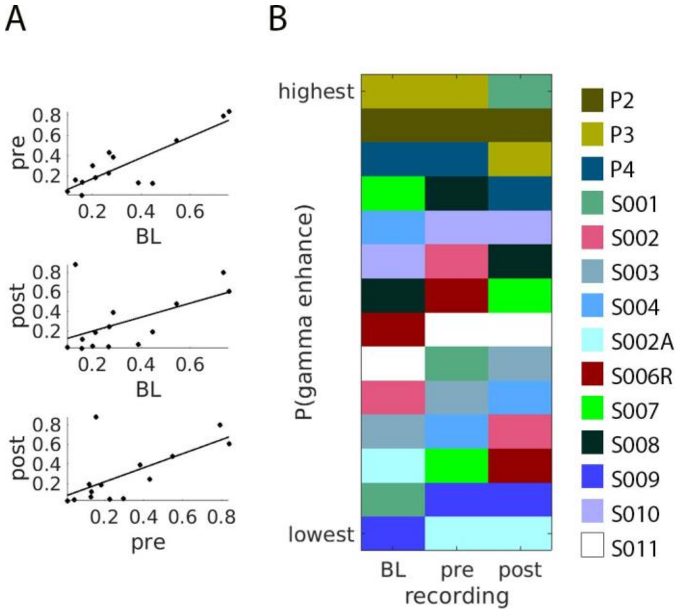


**Fig. S6**: ***Robustness of the individual preference for the gamma-enhancing network state.*** The probability of visiting the gamma-enhancing network state was inferred from different recording sessions (BL = task baseline, pre = rest recording preceding the task, post = rest recording following the task). A) Correlation between recording-specific estimates of time-average state probability. B) Ordering of subjects with respect to the probability of visiting the gamma-enhancing rest state. P1 was omitted because this subject did not have resting-state recordings.

**Tab. 1:** Parcel information. Parcel labels and Montreal Neurological Institute (MNI) coordinates of parcel centroids in millimetres. Centroid coordinates were obtained by averaging the coordinates of all sources of the cortical grid assigned to the respective parcel (see Materials and Methods). R = right; L = left.

| parcel label | X | Y | Z |
|---|---|---|---|
| Posterior Cingulate R | 11.89 | -51.50 | 15.92 |
| Posterior Cingulate L | -11.12 | -50.91 | 16.57 |
| Anterior Cingulate R | 9.23 | 26.70 | 14.12 |
| Anterior Cingulate L | -7.11 | 27.88 | 14.43 |
| Fusiform Gyrus R | 41.34 | -40.53 | -19.48 |
| Fusiform Gyrus L | -39.49 | -46.03 | -18.28 |
| Inferior Occipital Gyrus R | 36.74 | -85.05 | -9.23 |
| Inferior Occipital Gyrus L | -31.48 | -88.30 | -10.20 |
| Inferior Temporal Gyrus R | 52.10 | -21.98 | -25.09 |
| Inferior Temporal Gyrus L | -52.50 | -25.13 | -22.70 |
| Insula R | 40.92 | -12.10 | 12.67 |
| Insula L | -40.09 | -10.57 | 13.20 |
| Lingual Gyrus R | 16.75 | -73.55 | -4.56 |
| Lingual Gyrus L | -14.34 | -77.21 | -5.19 |
| Middle Occipital Gyrus R | 35.01 | -80.93 | 2.14 |
| Middle Occipital Gyrus L | -33.48 | -82.78 | 3.49 |
| Middle Temporal Gyrus R | 50.89 | -36.44 | -3.08 |
| Middle Temporal Gyrus L | -50.23 | -36.44 | -3.38 |
| Superior Temporal Gyrus R | 49.58 | -18.29 | -1.18 |
| Superior Temporal Gyrus L | -48.92 | -17.38 | -0.49 |
| Superior Occipital Gyrus R | 33.76 | -82.05 | 25.75 |
| Superior Occipital Gyrus L | -30.79 | -82.75 | 25.87 |
| Inferior Frontal Gyrus R | 42.97 | 23.20 | 4.31 |
| Inferior Frontal Gyrus L | -43.59 | 22.67 | 8.12 |
| Cuneus R | 15.80 | -86.33 | 17.68 |
| Cuneus L | -11.50 | -87.32 | 15.73 |
| Angular Gyrus R | 42.56 | -65.80 | 32.84 |
| Angular Gyrus L | -40.18 | -67.60 | 32.58 |

| | | | |
|---|---|---|---|
| Supramarginal Gyrus R | 48.28 | -49.33 | 33.37 |
| Supramarginal Gyrus L | -48.98 | -51.56 | 30.19 |
| Cingulate Gyrus R | 8.54 | -14.98 | 37.32 |
| Cingulate Gyrus L | -8.43 | -16.72 | 37.14 |
| Inferior Parietal Lobule R | 47.65 | -42.97 | 39.86 |
| Inferior Parietal Lobule L | -46.17 | -43.19 | 40.42 |
| Precuneus R | 16.83 | -65.32 | 42.40 |
| Precuneus L | -16.11 | -65.22 | 42.33 |
| Superior Parietal Lobule R | 27.28 | -61.83 | 56.16 |
| Superior Parietal Lobule L | -24.40 | -62.55 | 55.63 |
| Middle Frontal Gyrus R | 33.82 | 27.48 | 25.58 |
| Middle Frontal Gyrus L | -33.04 | 27.25 | 27.59 |
| Paracentral Lobule R | 8.73 | -36.21 | 60.96 |
| Paracentral Lobule L | -9.90 | -34.07 | 58.52 |
| Postcentral Gyrus R | 39.09 | -31.93 | 50.06 |
| Postcentral Gyrus L | -40.81 | -29.56 | 47.47 |
| Precentral Gyrus R | 43.96 | -11.88 | 40.57 |
| Precentral Gyrus L | -40.40 | -14.17 | 45.59 |
| Superior Frontal Gyrus R | 18.69 | 37.04 | 32.18 |
| Superior Frontal Gyrus L | -16.05 | 39.52 | 28.41 |
| Medial Frontal Gyrus R | 10.70 | 28.26 | 28.05 |
| Medial Frontal Gyrus L | -8.28 | 22.81 | 35.31 |
